# Bidirectional transcription marks accessible chromatin and is not specific to enhancers

**DOI:** 10.1101/048629

**Authors:** Robert S. Young, Yatendra Kumar, Wendy A. Bickmore, Martin S. Taylor

## Abstract

Bidirectional transcription initiating at enhancers has been proposed to represent the signature of enhancer activity. Here we show that bidirectional transcription is a pervasive feature of all forms of accessible chromatin, including enhancers, promoters, CTCF-bound sites and other DNase hypersensitive regions. Transcription is less predictive for enhancer activity than epigenetic modifications such as H3K4me1 or the accessibility of DNA when measured in both enhancer assays and at endogenous loci. Bidirectional transcription initiation from accessible chromatin is therefore not sufficient for, nor specific to, enhancer activity. The stability of enhancer initiated transcripts does not influence measures of enhancer activity and we cannot detect any evidence of purifying selection on the resulting enhancer RNAs within the human population. Our results suggest that transcription initiating at enhancers is frequently a by-product of promiscuous RNA polymerase activity at accessible chromatin, and may not generally play a functional role in enhancer activity.

---

Enhancers are modular, regulatory DNA elements that positively drive gene expression at a distance^1^. They are thought to be central to controlling cellular differentiation and developmental gene expression profiles, and mutations disrupting them have been associated with several Mendelian disorders^2, 3^. Widespread bidirectional transcription initiating proximal to enhancers has been observed^4, 5^ where the production of these enhancer RNAs (eRNAs) has been demonstrated to mark active enhancers^6^ and is correlated with increased expression from nearby, presumptive target promoters^7, 8^.

While most existing enhancer discovery methods are based on a characteristic chromatin profile (high H3K4me1 and low H3K4me3), this transcriptional signal has been advocated as a complementary approach^9^ and raises the intriguing possibility that enhancer RNAs themselves, or the action of transcription is mechanistically important for enhancer activity. Of candidate enhancers defined solely using RNA-seq evidence in mouse embryos, only 42% were validated using transgenic assays^6^, while the FANTOM5 consortium used Cap Analysis of Gene Expression (CAGE) transcriptome data and validated 67-74% of their predictions^9^. However, these validation rates are lower than the 75% obtained when enhancers are defined by their chromatin marks alone^10^ and is also lower than the 87% validation rate for enhancers defined by the binding of histone acetyltransferase p300^11^. It remains to be seen whether epigenetic marks or bidirectional transcription is more specific and accurate in identifying active enhancers.

The potential functionality of some of these eRNAs has been tested experimentally. siRNA knockdowns of a number of candidate eRNAs have resulted in reduced gene expression^12, 13^. Others have tethered the eRNA molecule to its cognate enhancer and shown that the mature eRNA molecule is required for enhancer activity^14, 15^. Several eRNAs have also been reported to be responsible for RNA polymerase II recruitment at the target promoter^13, 16^. At the human growth hormone gene locus, however, it is only the act of transcription which is correlated with enhancer activity, the transcribed sequence can be replaced with no effect on resulting gene expression^17, 18^. An analysis of 124 mouse eRNAs detected no evolutionary constraint within their exonic sequences^8^ which suggests that these mature transcripts are not generally required for enhancer function. Despite this convincing evidence for functionality of a handful of eRNAs^19^, there is likely a reporting bias against those that do not show an effect and the majority of the thousands of eRNAs identified to date have yet to be experimentally interrogated.

In this study, we investigate the specificity and importance of bidirectional transcription for enhancer identification and function. We show that both stable and unstable bidirectional transcription initiation can be detected at open chromatin regions that are not marked as enhancers and do not exhibit enhancer activity. Bidirectional transcription alone cannot detect regulatory regions with increased enhancer activity, and measuring this is less specific than measuring chromatin accessibility when attempting to identify gene promoter targets. Furthermore, mature eRNAs do not show evidence for purifying selection within the human population which argues against a function for these transcripts. We propose that bidirectional transcription is a by-product of an opening of chromatin at all types of regulatory regions, and is not sufficient for identifying active enhancers within the genome.

## Results

We investigated the specificity of bidirectional transcription to predict enhancers in four well studied cell lines (Gm12878, HepG2, Huvec, and K562). Sites of transcription initiation were identified through CAGE and complementary data^20^ (GRO-cap, PRO-seq) that can be used to detect the transcription start sites of both stable and rapidly degraded transcripts (see Methods). We found that transcription initiates bidirectionally from DNase1 hypersensitivity sites (DHSs) within enhancer regions defined by chromatin marks consistent with earlier findings^4–6,9^. However, we also observed similar patterns of transcription initiation proximal to DHSs in CTCF-bound regions and at remaining DHSs which overlap heterogeneous ‘Other’ chromatin state annotations (Fig. 1, Supp. Fig 1–4, Supp. Table 1-2). This confirms that, while much bidirectional transcription at enhancers does not produce stable RNA transcripts^19^, frequent unstable transcription is not specific to enhancers and is similarly seen at non-enhancer DHSs.

**Figure 1:**
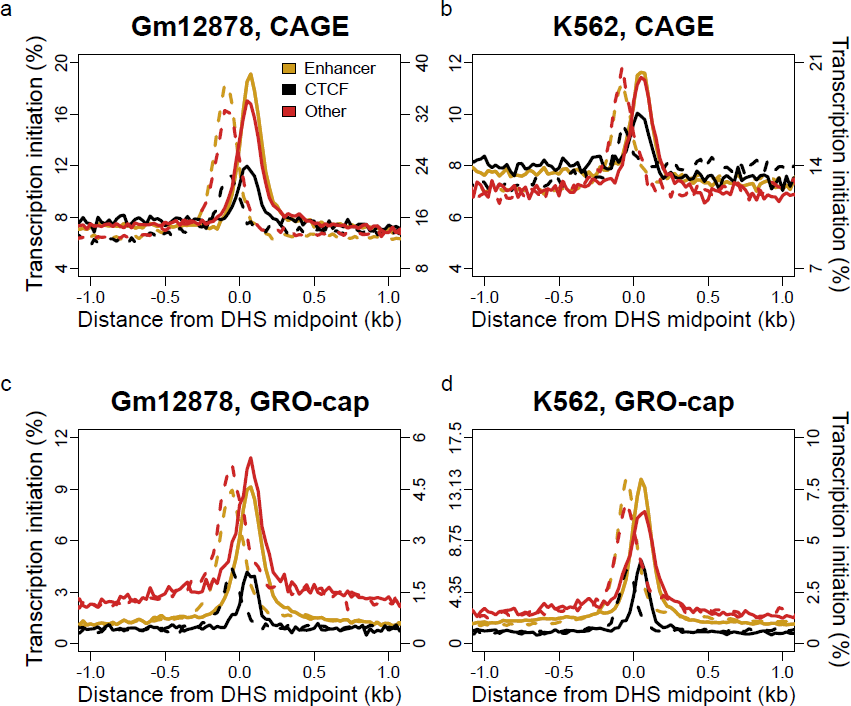
Bidirectional transcription initiates around DHSs but is not a specific mark of active enhancers. Stable bidirectional transcription across DHSs which do not overlap annotated promoters as measured by CAGE (top row) and unstable bidirectional transcription as measured by GRO-cap (bottom row) in Gm12878 and K562 cells. Solid lines consider transcription initiation from the positive strand and dashed lines show transcription initiation from the negative strand. The secondary axis on the top row corresponds to the level of transcription from the ‘Other’ DHSs which overlap a chromatin state annotation that is neither ‘Enhancer’ nor ‘CTCF’, while the secondary axis on the bottom row corresponds to the level of transcription from both the ‘Other’ and ‘CTCF’ DHSs.

**Supplementary Figure 1:**
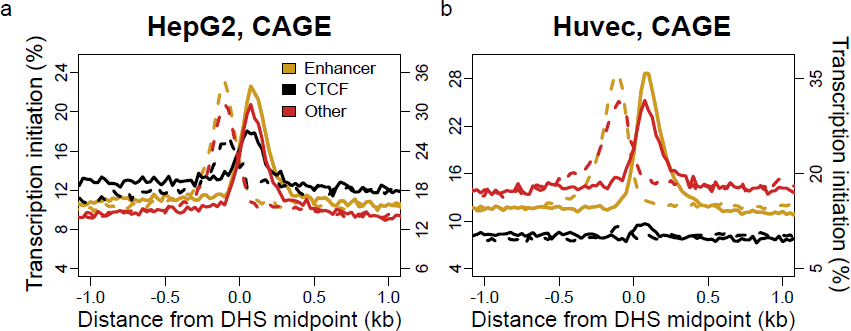
Bidirectional transcription initiates around DHSs but is not a specific mark of active enhancers. Stable bidirectional transcription across DHSs which do not overlap annotated promoters as measured by CAGE in HepG2 and Huvec cells. Solid lines consider transcription initiation from the positive strand and dashed lines show transcription initiation from the negative strand. The secondary axis on the top row corresponds to the level of transcription from the ‘Other’ DHSs which overlap a chromatin state annotation that is neither ‘Enhancer’ nor ‘CTCF’.

**Supplementary Figure 2:**
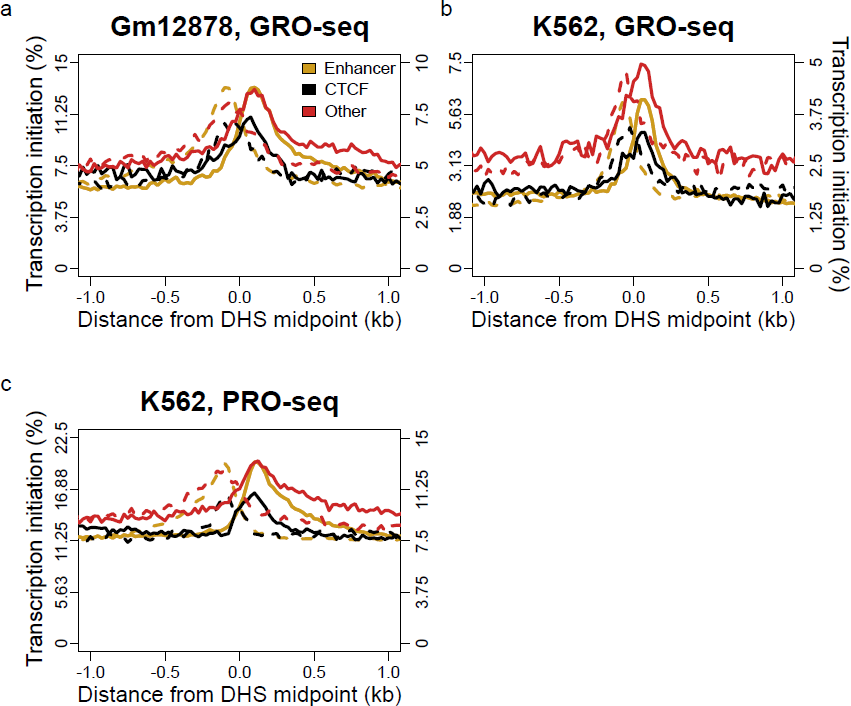
Bidirectional transcription initiates around DHSs but is not a specific mark of active enhancers. Unstable bidirectional transcription across DHSs which do not overlap annotated promoters as measured by GRO-seq in Gm12878 and K562 (a and b) cells and PRO-seq in K562 cells (c). Solid lines consider transcription initiation from the positive strand and dashed lines show transcription initiation from the negative strand. The secondary axes correspond to the level of transcription from the ‘Other’ and ‘CTCF’ DHSs which do not overlap the ‘Enhancer’ chromatin state annotation.

**Supplementary Figure 3:**
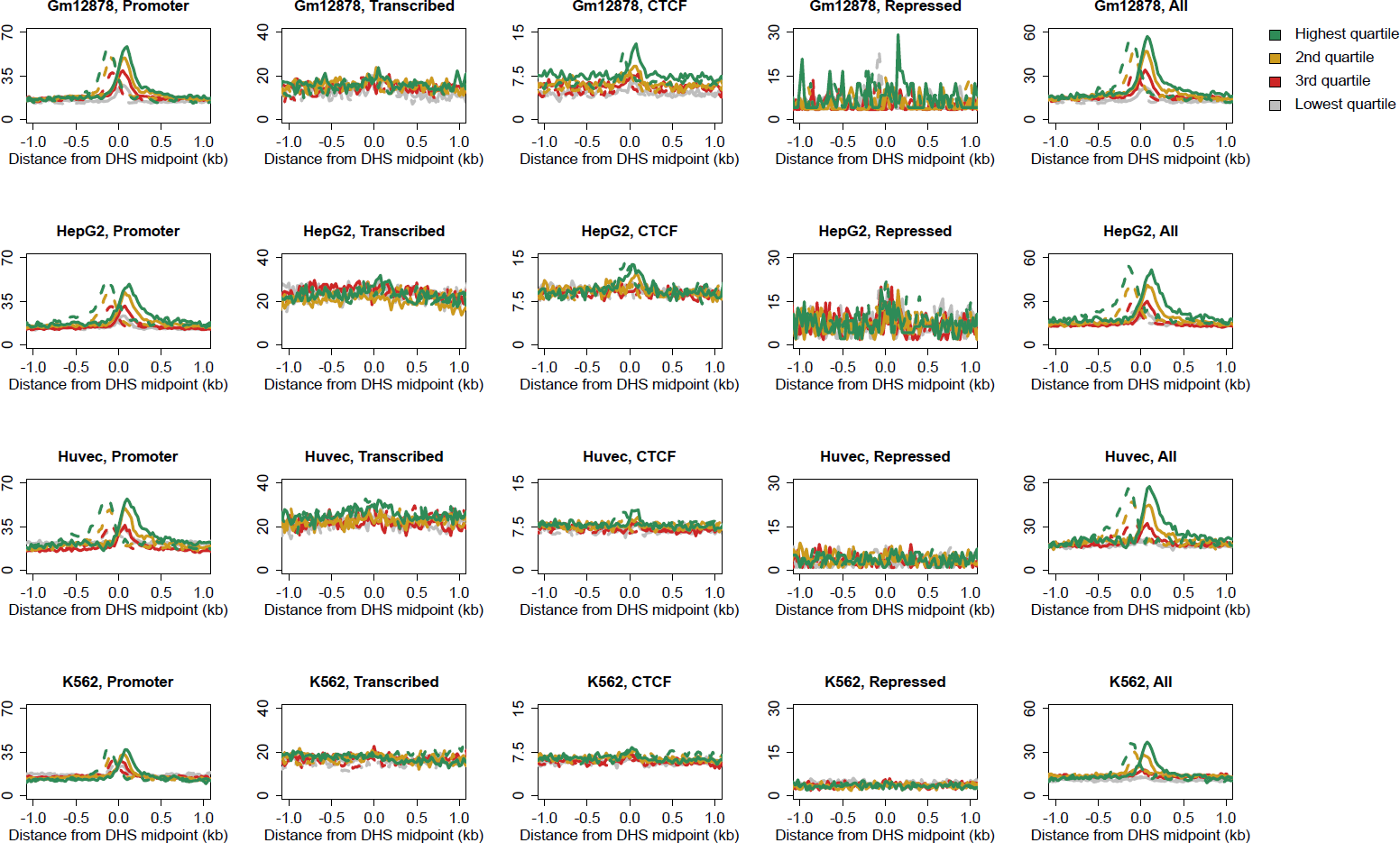
Stable bidirectional transcription initiation across DHSs which do not overlap annotated promoters in four cell lines across various chromatin state annotations. DHSs for each cell line are split into quartiles of increasing DHS peak height. The solid lines consider CAGE tags from the positive strand only and the dashed lines consider CAGE tags from the negative strand only.

**Supplementary Figure 4:**
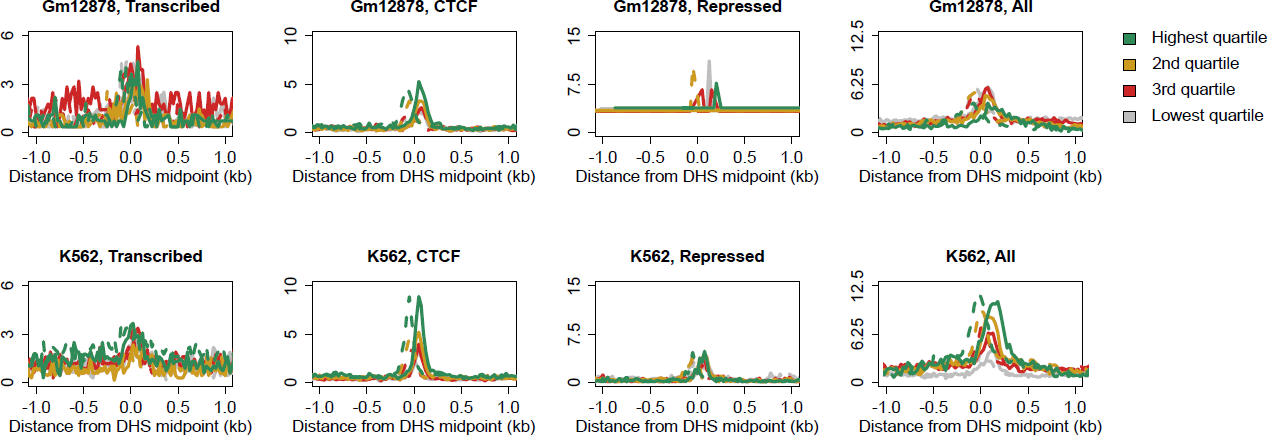
Unstable bidirectional transcription initiation across DHSs which do not overlap annotated promoters in four cell lines across various chromatin state annotations. DHSs for each cell line are split into quartiles of increasing DHS peak height. The solid lines consider GRO-cap tags from the positive strand only and the dashed lines consider GRO-cap tags from the negative strand only.

In all of these genomic settings, the fraction of DHSs with detected transcription was proportional to the strength of the DHS signal. This relationship can be detected when comparing DHS strength to the frequency of both stable and unstable transcription as measured by GRO-cap and CAGE (Supp. Fig. 3,4). This suggests that either the presence of accessible chromatin facilitates transcription initiation or, perhaps, that the act of transcription may itself be responsible for driving an increased chromatin accessibility. The nucleotide span of DHSs is largely consistent across these chromatin state annotations (Supp. Fig. 5), which also suggests that there could be a common mechanism linking DNA accessibility and transcription at enhancers and all other active regulatory elements within the genome. As none of these behaviours are specific to any chromatin state annotation studied here, we conclude that neither stable nor unstable bidirectional transcription initiation represent a specific mark for identifying enhancer elements.

**Supplementary Figure 5:**
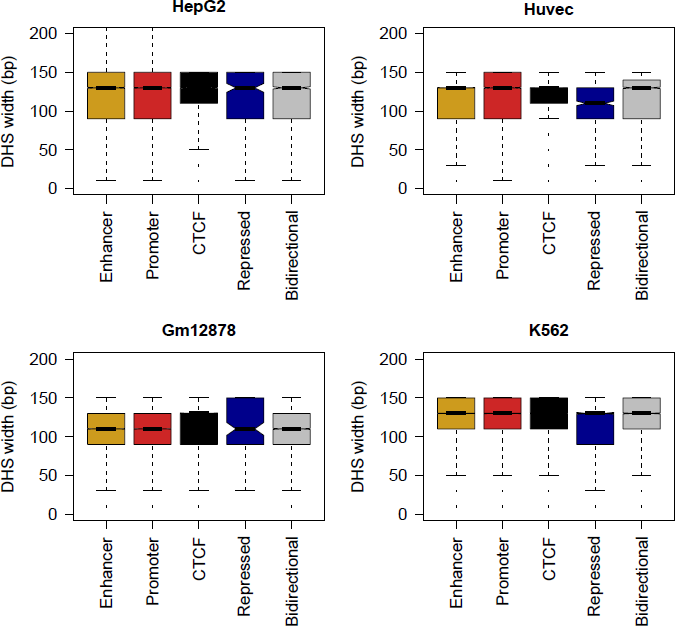
Widths of DNase hypersensitivity sites that overlap different chromatin state annotations and bidirectionally transcribed-defined enhancers across the four cell types studied here.

To further explore the relationship between transcription initiation and enhancer activity we intersected data from high-throughput enhancer reporter assays of candidate regulatory elements in K562 cells^21^ with CAGE and GRO-cap measures of transcription initiation at their endogenous genomic loci. Elements with transcription initiation and repressive chromatin marks do not exhibit enhancer activity relative to scrambled controls (median activity 0.9-fold, Mann-Whitney *p* = 0.01), demonstrating that neither bidirectional nor unidirectional initiation of transcription alone predicts enhancer activity (Fig. 2a). In contrast, histone modification-based chromatin state assignments do predict enhancer activity relative to scrambled controls (median activity 1.2-fold, Mann-Whitney *p* < 2.2×10^−16^), and a significantly greater measured activity for those elements marked as enhancers than those regions with repressive chromatin marks (median 1.3-fold, Mann-Whitney *p* < 2.2×10^−16^)^21^. Enhancers producing stable transcripts were not significantly more active in the reporter assays than those producing only unstable transcripts (Mann-Whitney *p* = 0.5) suggesting that neither transcript stability nor the transcripts themselves are generally required for enhancer activity. However, the subset of chromatin-defined enhancers without detected transcription initiation showed significantly lower reporter activity than the transcribed enhancers (median 0.8-fold, Mann-Whitney *p* = 0.01) albeit with a suggestively higher median activity (1.1-fold, Mann-Whitney *p* = 0.2) than all categories of elements with repressive marks (Fig. 2a). This relationship could be interpreted as being causal, where transcription boosts enhancer activity, or consequential with more active enhancers being more accessible to RNA polymerase.

**Figure 2:**
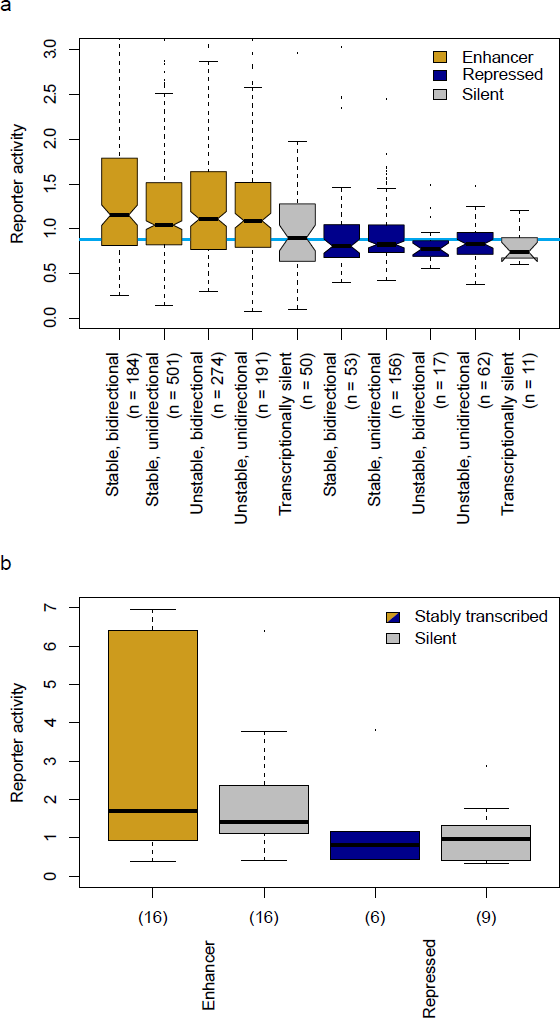
Transcription is not indicative of enhancer activity. (a) Reporter activities for enhancer and repressed regions in K562 cells with stable and unstable bidirectional and unidirectional transcription initiation, and those with no evidence for transcription. The blue line indicates the median reporter activity for all scrambled control sequences assays in K562 cells. (b) As for a, in HepG2 cells but only considering transcribed regions to be those with stable, bidirectional transcription initiation.

To further test our observation that bidirectional transcriptional initiation from accessible chromatin is not specifically associated with enhancer activity we performed our own additional reporter assays in HepG2 cells (Fig. 2b), These experiments were performed on enhancer regions specific to HepG2 cells, which are therefore not present in our above analyses of K562 enhancers. Again these results showed that chromatin marks effectively discriminate enhancers from repressed regions (median 1.7-fold greater reporter activity at enhancers, Mann-Whitney *p* = 0.01) but that there is only a minimal relationship between detectable bidirectional transcription and enhancer activity once conditioned on chromatin marks (Mann-Whitney *p* ≥ 0.47).

As enhancers are defined by their ability to positively drive gene expression in *cis*^1^, we next investigated the correlation between proposed markers of enhancer activity and transcription initiation from the closest annotated genic promoter. The correlation was carried out across the four well studied cell lines where matched chromatin state map, DHS and CAGE data were all available. To avoid the confounding influence of overlapping gene transcription we only considered candidate regulatory sites that were not contained within the extent of annotated genes nor within 1 kb of their boundaries (Fig. 3a). We find that regardless of chromatin state, typically 6 to 7% of candidate regulatory elements show (nominally significant) positively correlated transcription initiation with transcription initiation at the nearest genic promoter (Fig. 3b). Enhancers do not show a markedly increased frequency of correlation relative to CTCF or sites with repressive chromatin marks and are modestly less correlated that intergenic sites that exhibit chromatin marks characteristic of promoter activity (orphan promoters). If we consider candidate regulatory elements defined as previously advocated^9^ solely on the basis of bidirectional transcription initiation we again get the same approximately 7% fraction of positively correlated with the presumptive target (Fig. 3b). This result is consistent with previous observations of correlated expression between adjacent transcriptional units^22^, regardless of the function of these adjacent sites of transcription initiation, and suggests that this is driven by regional changes in transcriptional activity over a locus rather than defining the activity of discrete functional elements such as enhancers.

**Figure 3:**
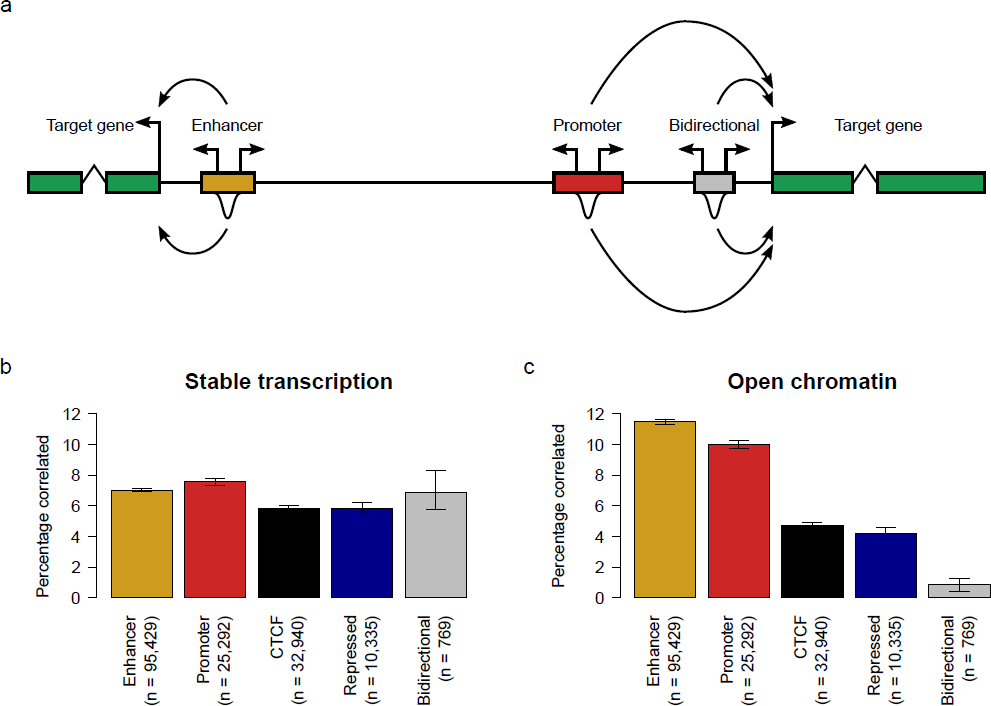
Stable transcription is not indicative of enhancer activity. (a) As shown by the curved arrows, the putative target of each chromatin state locus and bidirectionally transcribed-defined enhancer is defined as the nearest annotated gene (shown in the green boxes).The activity of each locus as measured by either the level of transcription initiation (the bidirectional arrows above each regulatory region) or the strength of the DHS signal (the peaks below each regulatory region) is then correlated with transcription initiation at the putative target gene promoter. (b) The percentage of chromatin state loci and bidirectionally transcribed-defined enhancers whose measure of stable transcription initiation is significantly correlated with transcription initiation from the nearest annotated gene promoter. The error bars represent the 95% confidence interval from 1,000 samplings of the data with replacement, while the numbers below each bar denote the number of loci tested for a significant correlation. (c) As for b, but the correlations being considered are between the level of DHS signal of chromatin state loci and bidirectionally transcribed-defined enhancers and transcription initiation from the nearest annotated gene promoter.

In stark contrast to transcription-based correlations between regulatory elements and genic promoters, DNase hypersensitivity measures do show clear discrimination between chromatin states in their correlation with genic transcription (Fig. 3c). DNase hypersensitivity at enhancer-marked regions is better correlated with transcription of the nearest gene than hypersensitivity associated with any of the other chromatin state categories (Fig. 3c). Enhancer hypersensitivity appears to have both greater sensitivity (Supp. Table 3; 10,961/6,713 = 63% more sites identified) and specificity (11.5% *vs.* 7.0%) than enhancer transcription for the identification of regulatory correlation (Fig. 3b,c). Defining candidate enhancers based solely on bidirectional transcription from a limited number of cell types (n = 4) identifies a relatively small number of sites whose transcription is no better correlated with genic expression than other intergenic genomic sites (Fig. 3b)

These results are robust as to whether genic expression was measured as the highest level of transcription from a single TSS (Fig. 3), or as the sum of CAGE tags over all annotated promoters for each protein-coding gene (Supp. Fig. 6). It is therefore the level of the open chromatin (as measured by DHS signal strength) at these sites, and not their transcriptional output, which can best be used to specifically identify enhancers and then associate them with putative promoter target(s).

**Supplementary Figure 6:**
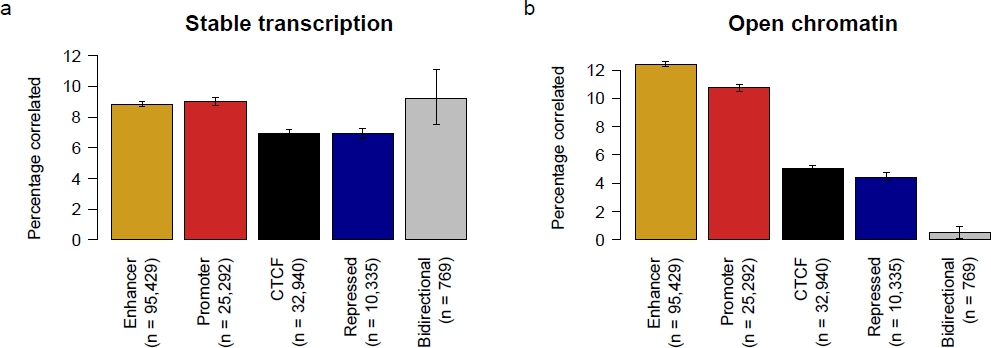
(a) The percentage of chromatin state loci and bidirectionally transcribed-defined enhancers whose measure of stable transcription initation is significantly correlated with the summed transcription initiation across the nearest annotated gene promoter. The error bars represent the 95% confidence interval from 1,000 samplings of the data with replacement, while the numbers above below bar denote the number of loci tested for a significant correlation. (b) As for a, but the correlations being considered are between the level of DHS signal of chromatin state loci and bidirectionally transcribed-defined enhancers and the summed transcription initiation across the nearest annotated gene promoter.

Having found that the stability of eRNA transcripts does not correspond to measures of enhancer activity (Fig. 2a) we took a complementary approach to test for organism level biological function in eRNAs by looking for evidence of evolutionary conservation or selective pressures on the DNA sequences encoding these molecules. If the mature eRNA is the functional moiety, we would expect these signals to be concentrated within their exonic, rather than intronic, sequence. This approach requires us to limit analyses to multi-exonic transcripts which must by definition be identified by RNA-seq (see Methods) rather than with CAGE or GRO-cap data. Consistent with a previous observation of mouse eRNA loci^8^, we did not detect any significant evolutionary constraint within eRNA exons when aligned between human and mouse (Supp. Fig. 7). However, the rapid gain and loss of non-coding regulatory elements through evolution^23, 24^ could potentially mask lineage specific functional constraint when considering deep (between species) sequence comparisons.

**Supplementary Figure 7:**
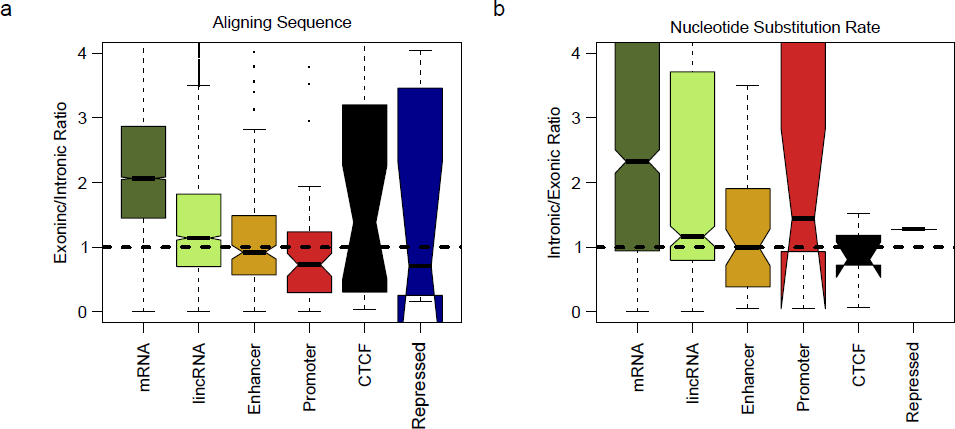
(a) Exonic/intronic ratio of the percentage aligning sequence between human and mouse for transcripts overlapping different genome annotations. The dashed line at one indicates equal conservation in exonic and intronic sequence. (b) Exonic/intronic ratio of nucleotide substitution rates of aligning sequence between human and mouse for transcripts overlapping different genome annotations. The dashed line at one indicates equal constraint in exonic and intronic sequence.

Measuring selective constraint within the human population, we compared the frequency of rare *vs.* common derived allele frequencies^24^ in exonic *vs.* intronic sequence. Purifying selection would be indicated by a relative excess of rare alleles in exonic sequence and positive selection indicated by a corresponding depletion of rare alleles. As expected, we observed strong purifying selection within mRNA exons, but no evidence of purifying selection in eRNA exons (Fig. 4). Similarly we did not see evidence of purifying selection in lincRNA exons consistent with previous reports^25^. In contrast to the case for eRNA and lincRNA, there is evidence for purifying selection within transcripts initiating proximally to intergenic promoter and, to our surprise, CTCF marks (Fig. 4). These results suggest that there is no widespread purifying selection at eRNA exonic sequences – either between species or within the human population – and again suggests that it is unlikely that the majority of mature eRNA transcripts examined here are biologically functional.

**Figure 4:**
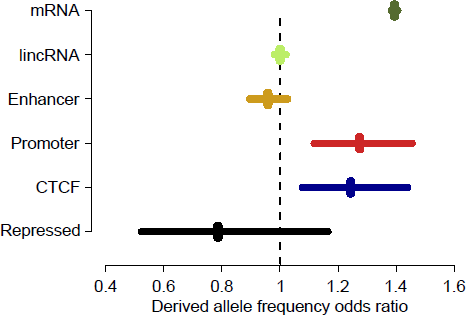
Mature eRNAs do not show evolutionary signatures of functionality. Odds ratios of derived allele frequencies for rare (< 1.5%) and non-rare (> 5%) derived alleles compared between exonic and intronic sequences for transcripts overlapping different genome annotations in the DeCode population. Horizontal lines indicate the 95% confidence interval of the odds ratio estimates. Odds ratios significantly greater than 1 indicate increased selective constraint in exonic relative to intronic sequence.

## Discussion

We have shown that low levels of transcription initiation are a common feature of accessible chromatin sites both associated with enhancer activity and those with other functions (Fig. 1,2). Furthermore, increased accessibility of the site (as measured by DNase hypersensitivity), is associated with an increased probability of detecting those transcription initiation sites (Supp. Fig. 3,4). This is consistent with a model of promiscuous RNA polymerase II transcription initiation on accessible DNA^26^ and a major role for chromatin in suppressing spurious transcription initiation^27^. DNA accessibility is not the sole determinant of transcription initiation, as nuclear position and the presence of specific transcription factor transactivation domains can dramatically influence transcription output^28^. However, it seems likely that the pervasive, low-level initiation of transcription associated with all categories of highly accessible chromatin represents a form of biological noise rather than specific activity required for the regulation of gene expression. It is clear from the data presented here that the bidirectional initiation of transcription at enhancers is not sufficient to elicit enhancer activity (Fig. 2) nor is bidirectional initiation specific to enhancer activity (Fig. 1).

Despite the lack of specificity for enhancers, measuring bidirectional transcription initiation is certainly not without merit, as it can be used to identify regions of open chromatin in exactly the same sample and source data in which gene expression is quantified^6, 9, 29^. The apparent success of bidirectional transcription alone in defining active enhancers (~70% validation rate^9^) can be explained by the observation that the majority of DNase hypersensitive, and thus transcription-initiating regions outside of genes, are in the context of chromatin defined-enhancers (Fig. 3, Supp Table 3). Transcription initiation provides positive predictive value for accessible DNA, but no power to discriminate enhancer from non-enhancer. Further, the measured transcriptional activity of enhancers is no more correlated with putative target gene expression than other categories of highly accessible DNA and is substantially less well correlated than DNase hypersensitivity in the context of enhancer chromatin marks (Fig. 3).

The pervasive low level initiation of transcription at highly accessible chromatin in diverse contexts suggests the resultant transcripts may be by-products rather than functional entities. There is an important distinction to be made between a molecular measure of function where there is a detectable molecular species or event; and a biological measure of function where the molecular species or event impacts an organism level phenotype. With current technologies we have the power to very sensitively detect the molecular products of the genome (<0.002 copies per cell for the CAGE libraires used in this study^30, 31^), but are all of those products really consequential for the biology of the organism? Our measures of selection tell us that both in comparisons between mammalian species and within humans, the nucleotide sequence of transcripts initiating in enhancers is indistinguishable from expectation under neutral evolution (Fig. 4, Supp. Fig. 7). This does not rule out the possibility that a minority of such sequences are important for organism biology, but overwhelmingly their sequence appears inconsequential for survival or reproductive fitness.

The observation that chromatin marked enhancers work equally well as enhancers whether their associated eRNAs are relatively stable or rapidly degraded (Fig. 2a), supports our measures of selective constraint in suggesting that eRNAs are not generally functionally important products. However, we cannot exclude the possibility that the action of transcription at enhancers (and other DHS), rather than the resultant transcript, is important for function or maintaining regulation at the site. Indeed, finding that chromatin marked enhancers without any detected transcription tend to exhibit lower enhancer activity than those with stable or unstable transcription (Fig. 2a) may support this view. Directly testing this possibility is challenging as any perturbation of transcription initiation, for example by targeting a transcriptional repressor to an enhancer, could also directly suppress the target promoter through looping or nuclear domain interactions^32^.

We propose that bidirectional transcription is predominantly a by-product of an opening of chromatin at all types of regulatory regions and, notwithstanding those published examples of functional eRNAs^19^, the majority of the transcripts produced are not likely to be required for regulatory function.

## Methods

### Genome annotation

Protein-coding, miRNA and lincRNA annotations were extracted from the GENCODE v14 release (June 2012). The promoters for these transcripts were recorded as −300/+100 bp around their annotated TSSs.

Chromatin state maps produced by the SEGWAY algorithm^33^ were downloaded for Gm12878, HepG2, Huvec and K562 cells from the Ensembl Biomart site (Release 67, May 2012)^34^. The states ‘Predicted Enhancer’ and ‘Predicted Weak Enhancer/Cis-reg element’ were merged into a single ‘enhancer’ state while the states ‘Predicted Promoter with TSS’ and ‘Predicted Promoter Flank’ were merged into a single ‘promoter’ state. For our cell-specific analyses, the ‘Other’ category includes all chromatin state annotations that do not overlap an ‘enhancer’ state or any CTCF-binding locations (defined below). A unified state map was built by merging each state annotation across cell types and then annotating the genome with the merged annotations using the following hierarchy: (1) enhancer, (2) promoter, (3) transcribed, (4) CTCF, (5) repressed. The transcribed regions marked in this manner were not considered in subsequent analyses. In this way, for example, a region is marked as an enhancer if it is annotated as such in at least one of the four cell types but a region is only annotated as repressed if it is marked as repressed in at least one cell type and is also not annotated by any of the other states in any cell type. Bidirectional transcription was measured only at annotated regions over 1 kb from an annotated GENCODE or RefSeq promoter while to further remove any confounding effects of neighbouring gene expression only regions over 1 kb from annotated GENCODE or RefSeq gene models were considered in the correlation analyses.

The genomic spans of bidirectionally transcribed-defined enhancers were obtained from http://enhancer.binf.ku.dk/presets/permissive_enhancers.bed^9^. As these enhancer predictions were defined using CAGE libraries from a wide range of cell lines and tissues, we filtered these to include only those loci which showed bidirectional transcription (defined by at least one overlapping CAGE tag on both the positive and negative DNA strand) in at least one of the four cell types considered here and which would be considered to be an active enhancer in at least one of the cell types by these authors. As for the chromatin state loci, enhancer predictions less than 1 kb from annotated GENCODE or RefSeq gene models were removed before performing the correlation analyses.

DNase1 hypersensitivity sites for each cell type^35^ were obtained directly from the UCSC genome browser (http://hgdownload.cse.ucsc.edu/goldenPath/hg19/encodeDCC/wgEncodeUwDnase/). We downloaded the ‘narrowPeak’ files for each cell type and considered only the intersection of both replicates in our analyses.

The reads for CTCF ChIP-seq experiments produced by the ENCODE consortium^10^ were downloaded from the Gene Expression Omnibus (GSE26320). Reads were then mapped to the hg19 genome using bowtie^36^ where only uniquely-mapping reads were retained (using option -m 1). All other parameters were left at their default values. Regions significantly enriched in CTCF relative to the whole cell extract samples were identified using macs14^37^ where a maximum of two reads at each individual position were retained (option ‐‐keep-dup 2). All other parameters were left at their default values.

### Transcriptome analysis

CAGE data produced by the FANTOM5 consortium^30^ were downloaded in BAM format from http://hgdownload.cse.ucsc.edu/goldenPath/hg19/encodeDCC/wgEncodeRikenCage/ and all libraries from each cell type were then merged into a single BAM file. GRO-cap, GRO-seq and PRO-seq data for K562 and Gm12878 cells^20^ were obtained from the GSE60456 series at the Gene Expression Omnibus (GSM1480321, GSM1480323, GSM1480325, GSM1480326, GSM1480237). Unstably-transcribed chromatin state annotations were identified as those with transcription initiation supported by GRO-cap, GRO-seq or PRO-seq evidence and where the genomic extent of overlapping DHSs do not overlap any evidence for stable transcription initiation as measured by CAGE.

The expression level for individual annotated regions across the unified chromatin state map was quantified for each cell type as the number of reads per kilobase region per million reads mapped (RPKM) summed across all libraries from that cell and also as the maximum RPKM from an individual TSSs location for each region. The expression of annotated GENCODE promoters (see above) were quantified in the same way.

Mapped RNA-seq reads from the ENCODE project^38^ were downloaded from the UCSC genome browser (http://hgdownload.cse.ucsc.edu/goldenPath/hg19/encodeDCC/wgEncodeCshlLongRnaSeq/) and we then assembled individual sequencing runs into transcripts using Cufflinks^39^, where the GENCODE v14 gene models were supplied as a guide reference annotation (option ‐g). All other parameters were left at their default values. All transcripts from all cell types were merged into a single set using the Cuffcompare program. Transcript expression was quantified across cell types and subcellular fractions as the number of fragments per kilobase exon per million reads mapped (FPKM) using Cuffdiff, which was run separately for each cell type. In order to control for genomic contamination, only those loci which contained at least one multi-exonic transcript or a single-exonic transcript with an FPKM > 1 in at least one subcellular fraction were considered for subsequent analyses.

### Linear correlations

We determined the closest annotated promoter to each chromatin state region and bidirectionally transcribed-defined enhancer and then calculated the linear correlation between transcription from that promoter and the transcription (as scored by CAGE RPKM) or strength of accessible chromatin (as scored by the DHS RPKM) at that chromatin state or bidirectionally transcribed-defined enhancer. If there were multiple, equally-close promoters to a given region then the correlation which gave the lowest *p*-value was considered. A positive correlation was recorded if the correlation coefficient was greater than 0 and *p* < 0.05. All other regions where considered to be nonsignificant. The uncertainty in the estimate of the percentage of positive correlations was determined by 1,000 samplings of the data with replacement.

### Reporter assays

Enhancer and repressor element activity in K562 cells was estimated as the mean expression values obtained from a parallel reporter assay^21^. For this analysis, stably-transcribed enhancers and repressed elements were identified as those with any evidence of stable transcription using CAGE originating from the entire chromatin state locus, regardless of DHS overlap. Unstably-transcribed elements are defined as those with no CAGE support over the locus, but evidence of transcription from at least one of the GRO-cap, GRO-seq, or PRO-seq datasets. The expression of sequence-scrambled controls for enhancer and repressor elements were not significantly different from each other (Mann-Whitney *p* = 0.20) and therefore these two categories were merged and considered as a single null expectation for the reporter activity measured from random DNA sequences.

Additional validations were performed in HepG2 cells (confirmed free of mycoplasma contamination with the Lonza MycoAlert kit). HepG2 cells were sourced from the Institute of Genetics and Molecular Medicine (Edinburgh) technical services. For our HepG2 reporter assays, DHSs from each chromatin state category which were more than 1 kb beyond RepeatMasker-marked regions and GENCODE gene annotations were randomly selected. PCR primers (Supp. Table 4) with Kpn1 and EcoRV sites were used to amplify 500-1500 bp regions containing the DHS site from HepG2 genomic DNA. Amplicons were cloned in to pGL4.26 vector post restriction digest. For reporter assays, pGL4.26 constructs and the pRLTK plasmid were co-transfected into HepG2 cells with Lipofectamine-2000. 48 hours post-transfection, firefly and renilla luciferase activity was measured from three replicates using the Promega dual luciferase kit. The firefly luciferase signal was normalized by the renilla luciferase signal to reduce variability in transfection efficiency and an average reporter activity for the three replicates was then calculated relative to the empty pGL4.26 vector.

### Evolutionary analysis

We extracted aligning sequence between human (hg19) and mouse (mm9) genomes from the 12-way mammalian EPO alignments (May 2012 release) from Ensembl^34^. Conservation was scored as the percentage of bases within a region which could be aligned between these two species. When estimating the substitution rate of the aligning material, we first removed the bases immediately adjacent to any alignment gaps, as these have previously been shown to bias substitution rate estimations^40^. Substitution rates were estimated using the HKY85 substitution model within the PAML package^41^, with those alignments giving a substitution rate greater than 10 were removed from subsequent analyses.

Derived allele frequencies were extracted from the deCODE large-scale whole-genome sequencing study of the Icelandic population^43^. Polymorphic SNPs were partitioned into rare (< 1.5% population frequency) and common (> 5% frequency) as previously^24^ and the frequency of these was compared between exonic and intronic sequence from different transcript types using Fisher’s exact tests.

## Acknowledgements

R.S.Y. and M.S.T. acknowledge the support of the UK Medical Research Council (MC_PC_U127597124) and the Medical Research Foundation. W.A.B. and Y.K. acknowledge the support of the UK Medical Research Council.

## Author Contributions

R.S.Y., W.A.B. and M.S.T. conceived and designed the experiments; R.S.Y. and Y.K. performed the experiments and analysed the data; R.S.Y. and M.S.T. wrote the paper.

## Competing Interests

The authors declare no competing financial interests.

